# Multi-channel 4D parametrized Atlas of Macro- and Microstructural Neonatal Brain Development

**DOI:** 10.1101/2021.02.11.430835

**Authors:** Alena Uus, Irina Grigorescu, Maximillian Pietsch, Dafnis Batalle, Daan Christiaens, Emer Hughes, Jana Hutter, Lucilio Cordero Grande, Anthony N. Price, Jacques-Donald Turnier, Mary A. Rutherford, Serena J. Counsell, Joseph V. Hajnal, A. David Edwards, Maria Deprez

## Abstract

Structural and diffusion MRI provide complimentary anatomical and microstructural characterization of early brain maturation. The existing models of the developing brain in time include only either structural or diffusion channels. Furthermore, there is a lack of tools for combined analysis of structural and diffusion MRI in the same reference space.

In this work we propose methodology to generate multi-channel (MC) continuous spatio-temporal parametrized atlas of brain development based on MC registration driven by both T2-weighted and orientation distribution functions (ODF) channels along with the Gompertz model (GM) fitting of the signals and spatial transformations in time. We construct a 4D MC atlas of neonatal brain development during 38 to 44 week PMA range from 170 normal term subjects from developing Human Connectomme Project. The resulting atlas consists of fourteen spatio-temporal microstructural indices and two parcellation maps delineating white matter tracts and neonatal transient structures. We demonstrate applicability of the atlas for quantitative region-specific comparison of 140 term and 40 preterm subjects scanned at the term-equivalent age. We show multi-parametric microstructural differences in multiple white matter regions, including the transient compartments. The atlas and software will be available after publication of the article.

## 1 INTRODUCTION

In addition to being a routine diagnostic tool in neonatal brain imaging (Rutherford et al., 2010), MRI has been widely used for quantification and interpretation of neonatal brain development in term- and preterm-born infants. Premature birth before 37 weeks postmenstrual age (PMA) is associated with an increased risk of atypical brain maturation leading to neurocognitive and neurobehavioural disorders. Multiple studies demonstrated correlation of MRI metrics with prematurity, clinical and environmental factors and neurodevelopmental outcomes (Ball et al., 2017; Barnett et al., 2018; Dimitrova et al., 2020). In this context, models of the normal brain development such as spatio-temporal atlases (Schuh et al., 2018) can also potentially facilitate detection of altered maturation patterns. The advanced acquisition and reconstruction protocols produce high-resolution structural T1-weighted (T1w) and T2-weighted (T2w) MRI volumes that allow segmentation of fine brain anatomical structures (Makropoulos et al., 2014). But these MRI modalities have low contrast for white matter (WM) structures that also varies during the neonatal stage due to the ongoing myelination. On the other hand, lower resolution diffusion MRI reflects the properties of tissue microstructural complexity in terms of diffusivity, anisotropy, neuronal density and fibre orientation (Bastiani et al., 2019; Pietsch et al., 2019). Combined diffusion and structural MRI analysis has already showed a potential to increase interpretability of the brain maturation patterns (Ball et al., 2017).

### 1.1 Structural MRI metrics

The structural MRI-derived metrics most commonly used in the neonatal brain studies include tissue- and structure-specific volumetry (Kuklisova-Murgasova et al., 2011; Makropoulos et al., 2016; Thompson et al., 2019) and surface measurements such as cortical thickness and curvature (Bozek et al., 2018; Fenchel et al., 2020) that can be extracted from automated segmentations (Makropoulos et al., 2014). Recently, automated segmentation of T2w images has been also applied for quantification of the volume of myelinated regions (Wang et al., 2019). Intensity changes in T1w and T2w images characterize white matter injury (O’Muircheartaigh et al., 2020) and diffuse excessive high signal intensity (DESHI) regions (Morel et al., 2020). Quantitative and semi-quantitative metrics applied to developing neonatal brain include the T1w/T2w signal ratio associated with myelin content (Bozek et al., 2018) and T2 relaxometry (Pannek et al., 2013; Kulikova et al., 2015; Wu et al., 2017; Knight et al., 2018).

### 1.2 Diffusion MRI metrics

Brain microstructure can be probed using a variety of quantitative metrics derived from diffusion MRI. Even though diffusion tensor imaging (DTI) is limited by inconsistencies in fiber-crossing regions (Jeurissen et al., 2013), DTI-derived metrics, including the fractional anisotropy (FA) and the mean, radial and axial diffusitivity (MD, RD and AD) are still most widely used in the neonatal brain studies (Barnett et al., 2018; Feng et al., 2019; Thompson et al., 2019; Dimitrova et al., 2020). Recently, higher order metrics, that alleviate some of the limitations of the DTI in the fibre crossing regions, have also been applied to investigate neonatal brain development, including the mean kurtosis (MK) index derived from diffusion kurtosis imaging (DKI) (Bastiani et al., 2019) and intracellular volume fraction (ICVF), fiber orientation dispersion index (ODI) and volume fraction of the isotropic compartment (FISO) derived from Neurite Orientation Dispersion and Density Imaging (NODDI) model (Zhang et al., 2012). The NODDI-derived indices have been used to characterize development of both white and gray matter microstructural features (Kunz et al., 2014; Batalle et al., 2019; Kimpton et al., 2020; Fenchel et al., 2020). The microscopic fractional anisotropy (*μ*FA) index (Kaden et al., 2016) designed to disentangle microscopic diffusion anisotropy from the orientation dispersion has not yet been applied to neonatal brain. Constrained spherical deconvolution (CSD) (Tournier et al., 2007) allows extraction of orientation-resolved microstructural information as orientation distribution functions (ODFs) from multi-shell high angular resolution diffusion imaging (HARDI) data. Based on fibre ODFs, fixel-based analysis (Raffelt et al., 2017) provides the means for assessment of specific fibre populations in terms of fibre density (FD) and fibre-bundle cross-section (FC) (Pannek et al., 2018; Pecheva et al., 2019).

### 1.3 Atlases and models of the neonatal brain development

Spatio-temporal normalisation and construction of age-specific group-average templates have been routinely employed in processing pipelines in the recent large neonatal brain MRI studies, to detect inter-group differences and anomalies in individual brains (Oishi et al., 2019). The majority of the reported spatio-temporal population-averaged atlases of the neonatal brain include either structural (T2w and T1w) (Kuklisova-Murgasova et al., 2011; Serag et al., 2012; Wright et al., 2014; Schuh et al., 2014; Makropoulos et al., 2016; Schwartz et al., 2016; Schuh et al., 2018; Wang et al., 2019; O’Muircheartaigh et al., 2020) or diffusion (Feng et al., 2019; Pietsch et al., 2019; Dimitrova et al., 2020) channels. Up to our knowledge, the only existing multi-channel population-averaged T1w+T2w+DTI atlas (Oishi et al., 2011) was constructed from a set of normal term subjects from 38 to 41 weeks PMA. However, the averaged template was reported to have significantly lower sharpness than the original T2w and DTI images. Apart from (Feng et al., 2019) who used FA+MD guided registration, these atlases were constructed using registration driven by a single channel and the output transformations were propagated to the rest. In general, one of the challenges of multi-channel registration is considered to be the alignment between the structural and diffusion MRI volumes. Following spatial normalization, the templates were created using either weighted or direct averaging of the signal in the reference space. Zhang et al. (2016) proposed to perform averaging in the frequency domain and reported higher sharpness of the atlas features.

Due to the rapid changes of structure, volume and cytoarchitecture during the fetal and neonatal period, the majority of the atlases have been also resolved in time in the form of weekly templates. Smooth transition between the atlas time points have been provided through kernel regression (Kuklisova-Murgasova et al., 2011; Serag et al., 2012; Schuh et al., 2014, 2018), logistic regression (Wang et al., 2019) or Gaussian process regression (Marquand et al., 2016; O’Muircheartaigh et al., 2020; Dimitrova et al., 2020). Recently, Gompertz model was successfully used to parametrize fetal and neonatal brain volumetry and surface measurements (Wright et al., 2014; Makropoulos et al., 2016; Schwartz et al., 2016), showing better approximation than the linear model (Makropoulos et al., 2016), even though the changes in averaged structural (O’Muircheartaigh et al., 2020) and DTI (Bastiani et al., 2019; Feng et al., 2019; Dimitrova et al., 2020) metrics in WM and GM can be approximated by linear trends. However, so far, there has been no reported works combining structural and diffusion MRI into a spatio-temporal atlas of the normal term born neonatal brain development.

### 1.4 Region specific analysis

The majority of the neonatal brain studies has been employing region-specific quantitative analysis based on correlation between the MRI-derived metrics measured within specific regions and parameters such as gestation age (GA) at birth, clinical factors or neurodevelopmental outcomes. In structural-only MRI datasets segmentation is normally performed by atlas-based methods (Makropoulos et al., 2014). In the WM atlas-based analysis, the parcellation maps for the single-subject or population-average WM DTI atlases (Oishi et al., 2011; Feng et al., 2019; Alexander et al., 2020) were created by 2D manual delineation based on DTI directionally-encoded colour maps for single subject or population-averaged templates. Label propagation based on DTI channel-guided registration has been widely used in neonatal brain studies (Kersbergen et al., 2014; Rose et al., 2014; Wu et al., 2017; Claessens et al., 2019; Feng et al., 2019). The tract-based spatial statistics (TBSS) (Smith et al., 2006) approach uses skeletonized FA maps for definition of the regions (Krishnan et al., 2016; Young et al., 2018; Barnett et al., 2018; Thompson et al., 2019). As an alternative, track-specific analysis employs tractography to identify and segment the major WM pathways (Kulikova et al., 2015; Akazawa et al., 2016; Pecheva et al., 2017; Bastiani et al., 2019; Zollei et al., 2019; Kimpton et al., 2020; Dubner et al., 2020). In this case, the seed regions for tractography are defined in the template space and the segmentations of WM tracts are achieved by thresholding of the resulting probabilistic tractography maps. In Akazawa et al. (2016), this approach was also used to create population-specific average probabilistic maps of the major WM tracts.

### 1.5 Contributions

In this paper we propose to merge multiple metrics extracted from both diffusion and structural MRI in a single parametrized multi-channel spatio-temporal atlas of the normal neonatal brain development.

The generated 4D MC atlas covers 37-44 weeks PMA range and includes structural (T1w, T2w and T1w/T2w myelin contrast) and diffusion channels with ODF, DTI, DKI, *μ*FA and NODDI derived metrics. Furthermore, the atlas includes two parcellation maps: (i) the major WM tract regions (Alexander et al., 2020) refined using probabilistic tractography in the template and (ii) the map of the transient WM regions with high maturation rates during neonatal period. To ensure accuracy of spatial alignment, we propose a multi-channel (MC) registration (Uus et al., 2020) guided by spatially-weighted structural, diffusion (ODFs) MRI and cortical segmentation (Makropoulos et al., 2018) channels. Parametrization in time is performed by the Gompertz model (GM) widely used for fitting of growth data. We implemented the atlas construction and fitting functionalities within MRtrix3 software package (Tournier et al., 2019). To demonstrate the application of the proposed atlas we perform multi-modality study to compare term and preterm brain development and identify regions where WM maturation has been altered by preterm birth.

## 2 MATERIAL AND METHODS

### 2.1 Cohort, datasets and preprocessing

The atlas was constructed using 170 multi-modal MRI datasets of term-born neonates (born and scanned between 37 and 44 weeks PMA) that included T1w, T2w and HARDI scans. Additional 40 datasets of preterm neonates (born between 23 and 32 weeks GA: 28.94∓2.54 and scanned between 37 and 44 weeks PMA) were used for comparison analysis. Inclusion criteria were high image quality for scans of all modalities, singleton pregnancies and no major brain abnormalities. All scans were acquired under the developing Human Connectome Project (dHCP)^1^.The distribution of the GA at birth and PMA at scan is given in Fig. 1.

**Figure 1.**
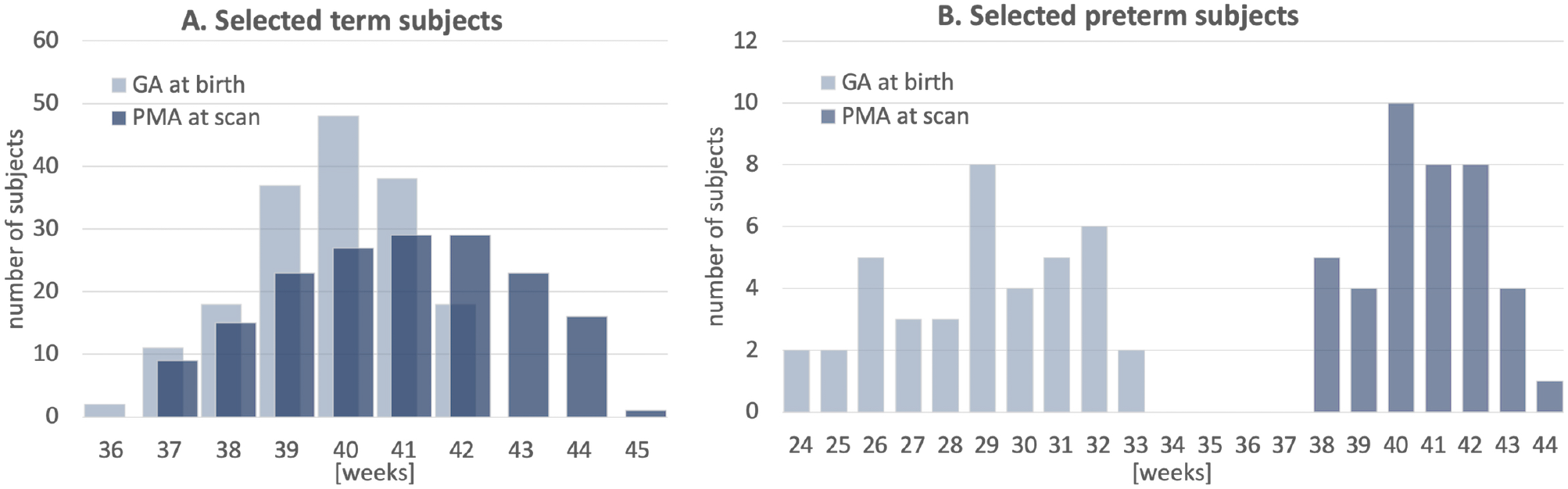
Selected cohort of neonatal subjects from dHCP project: GA at birth and PMA at scan of 170 term subjects **(A)** and 40 preterm subjects **(B)**.

Datasets were acquired on a 3T Philips Achieva scanner equipped with a dedicated 32-channel neonatal head coil and baby transportation system (Hughes et al., 2017). The multi-shell HARDI volumes were acquired with four phase-encode directions on four shells with b-values of 0, 400, 1000 and 2600 *s/mm*^2^ with TE 90 *ms*, TR 3800 *ms* (Hutter et al., 2018; Tournier et al., 2020) with 1.5 × 1.5 × 3 *mm* resolution and 1.5 *mm* slice overlap and reconstructed to 1.5 *mm* isotropic resolution using the SHARD pipeline (Christiaens et al., 2021). The structural T2w volumes were acquired using a TSE sequence with TR 12 *s*, TE 156 *ms*. The T1-weighted volumes were acquired using an IR TSE sequence with TR 4.8 *s*, TE 8.7 *ms*. The isotropic T2w and T1w volumes with 0.5 *mm* resolution were reconstructed using a combination of motion correction (Cordero-Grande et al., 2018) and super-resolution reconstruction (Kuklisova-Murgasova et al., 2012).

Intensities of individual T1w and T2w volumes were bias-corrected and normalised to the same intensity ranges. The brain tissue and structure segmentations were generated by dHCP pipeline (Makropoulos et al., 2014, 2018). For each dataset, the structural and diffusion volumes were coaligned using affine registration of T2w and MD volumes using NCC similarity metric in MRTrix3. The DWI volumes were globally normalised prior to the nonlinear multi-channel registration step (Tournier et al., 2019).

### 2.2 Extraction of MRI metrics

The structural metrics include normalised T1w and T2w intensities as well as the the T1w/T2w ratio thought to be associated with the myelin content (Glasser and Van Essen, 2011). Furthermore, we extracted Jacobians (J) of deformation fields from the MC registration output (Section 2.4) to measure local volumetric changes.

The DTI metrics included AD, MD, RD and FA extracted using MRtrix3 toolbox (Tournier et al., 2019). The DKI fitting and calculation of MK was performed similarly to Bastiani et al. (2019). The NODDI (Zhang et al., 2012) toolbox was used for fitting FISO, ICVF and ODI metrics. The estimation of micro FA maps was performed using SMT toolbox (Kaden et al., 2016). Only the two top HARDI shells were used for *μ*FA and DKI fitting in order to minimise the impact of artefacts. In addition, we computed the mean DWI signal mDWI for the top 2600 *s/mm*^2^ shell since it provides high contrast for WM structures. We extracted WM ODF from HARDI using MRtrix3 multi-shell multi-tissue constrained spherical deconvolution (CSD) (Jeurissen et al., 2014). The track density imaging (TDI) maps were generated from the outputs of probabilistic tractography (Tournier et al., 2010) with whole brain as the seed region and 700,000 streamlines for all dataset. This particular number of streamlines was selected arbitrary.

### 2.3 Multi-channel registration of structural and HARDI MRI

In this work, we propose a MC non-linear registration technique to improve accuracy of spatial normalisation. We build on multi-contrast ODF registration framework (Pietsch et al., 2017; Raffelt et al., 2011) which employs SyN Demons (Avants et al., 2007) with an SSD metric and reorientation of ODF using apodized point spread functions (Raffelt et al., 2012). In order to decrease the sensitivity to acquisition or physiology related changes in signal intensities, we propose to replace SSD metric with a novel robust local angular correlation metric for ODF channels based on angular correlation metric introduced in (Anderson, 2005). We further add structural and tissue parcellation channels with local NCC (LNCC) similarity measure. We combine the channels through weighted fusion of the displacement field updates (Forsberg et al., 2011).

Angular correlation *r_A_* between two ODFs *F^ODF^* and *G^ODF^* represented with real valued spherical harmonic (SH) orthonormal basis functions *Y_lm_*(*θ, ϕ*)

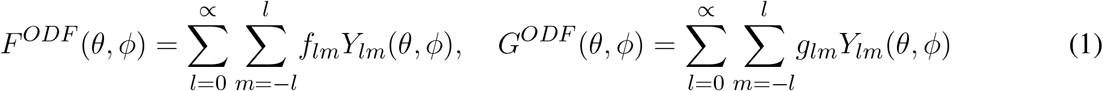

is computed as (Anderson, 2005):

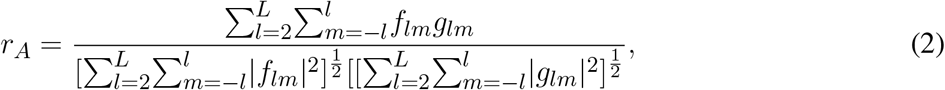

where *g_lm_* and *f_lm_* are the SH coefficients of *G^ODF^* (*θ, ϕ*) and *F^ODF^* (*θ, ϕ*) of order *L* with even *l* = {2, 4, …, *L*} harmonic degree terms, correspondingly. The *l* = 0 term does not contribute to AC values. Since this is a correlation metric, the corresponding symmetric updates to the displacement fields Λ^*F*^ and Λ^*G*^ can be computed in a similar manner to LNCC demons (Avants et al., 2008) but with respect to the 4D ODFs rather than only the 3D local neighbourhood (Eqn. 3).

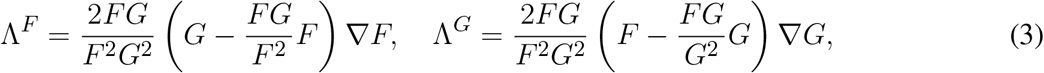

where 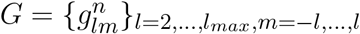 and 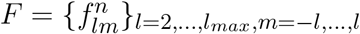 are the vectors of SH coefficients at a given location in the 3D volume space with local neighbourhood *n* = 1, …, *N*. We refer to this registration metric as local angular cross-correlation (LAC).

The contributions from each of the channels *i* to the global symmetric displacement field update Λ^*global*^ are locally weighted with respect to the 3D certainty maps based on the approach proposed in (Forsberg et al., 2011). First, at every iteration, the certainty gradient maps 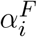 and 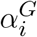 are computed from the original volumes *F* and *G* for each of the channels (including structural and ODF volumes) and normalised as:

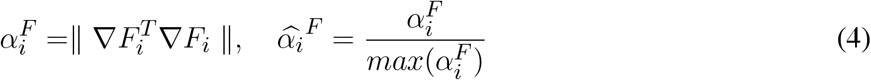

Then, the global symmetric updates to the displacement fields are computed by weighted averaging of the channel-specific update fields with respect to the certainty maps. The sum of all weighted updates is normalized with respect to the sum of the gradient maps.

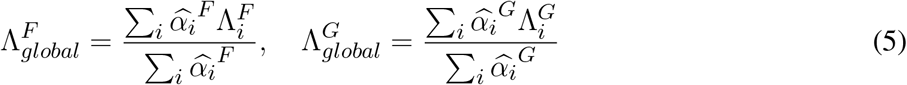

The proposed approach ensures that the output deformation fields are defined by the contribution of the local channel regions with the highest structural content. This is relevant for the ROIs where one of the channels has low intensity contrast. In comparison, the classical multi-variate SyN approach Avants et al. (2007) is based on averaging of the individual channel updates.

### 2.4 Generation of 4D multi-channel atlas

The 4D parametrized MC atlas of the neonatal brain development was generated from the 170 term neonatal datasets in three sequential steps: (A) initial registration of structural channels to a single structural template and creation of an average multi-channel template, (B) refined registration of structural and diffusion channels to the multi-channel template and creation age-dependent average multi-channel templates, (C) fitting of the signal and deformation fields in time using the Gompertz model to generate parametrized 4D multi-channel atlas. The proposed pipeline is summarised in Fig. 2.

**Figure 2.**
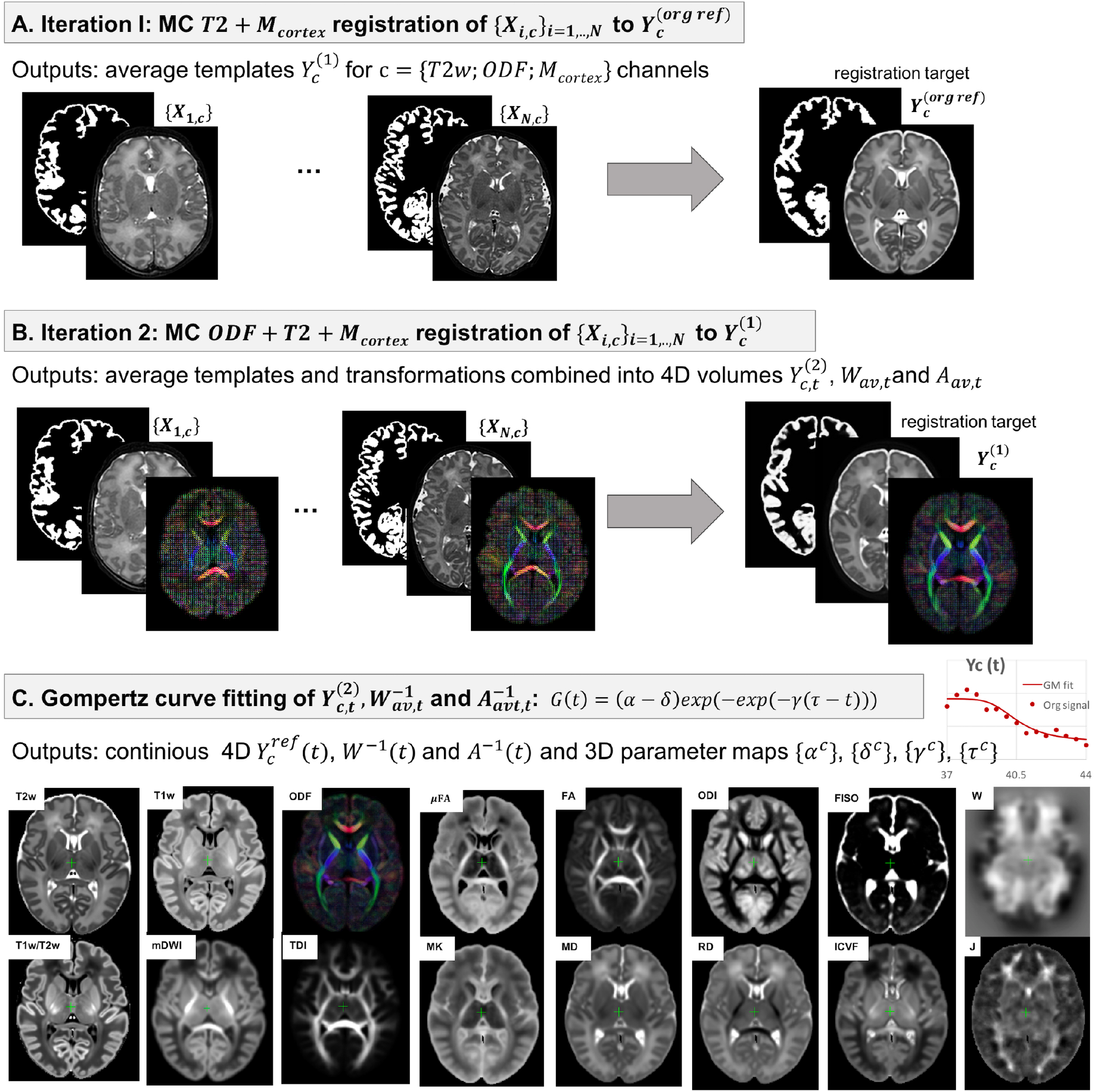
The proposed pipeline for generation of parametrized 4D MC atlas of the neonatal brain development during 37 to 44 weeks PMA range. **A:** (Iteration 1) The T2w + cortex mask guided MC registration to a global reference space *Y*^(*ref*)^ (36 weeks T2w dHCP template from Schuh et al. (2018)) is performed for all subjects. Preliminary average templates 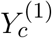 are created for T2w, cortex mask and ODF channels. **B:** (Iteration 2) The ODF + T2w + cortex mask guided MC registration to the new multi-channel template is performed of all subjects. Average templates 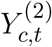 are created for all channels and 15 discrete time points. The average inverse nonlinear warps 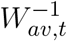 and affine transformations 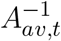 are also created for all 15 time points. **C:** The Gompertz curve fitting in time is performed for all channels and transformations resulting in continuous 4D multi-channel atlas 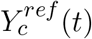 in the reference space the unbiased version *Y*_*c*_(*t*).

#### 2.4.1 Generation of a 3D multi-channel template

We chose the T2w 36 week template from the dHCP neonatal brain atlas^2^ (Schuh et al., 2018) as the global 3D reference space (*Y* ^(*reforg*)^) due to the lower degree of cortical folding which facilitates more accurate registration of the cortex. All datasets {*X_i_*},_*i*=1,…,*N*_ were registered to this template using affine alignment with global NCC followed by non-linear registration guided by two structural channels (T2w + cortex mask), similarly to O’Muircheartaigh et al. (2020):

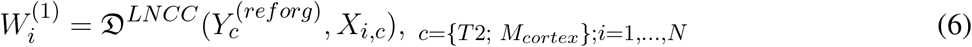

where 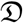 is the MC Demons registration operator, 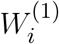 are the output deformation warps for each of the *N* datasets *X*_*i,c*_ with *c* = {*T*; *M*_*cortex*_} channels and 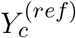 is the reference volume. The MC registration included spatially weighted fusion of the channels (Sec.2.3, Uus et al. (2020)). The output deformation warps 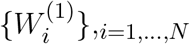 were propagated to the rest of the structural and dMRI channels. The preliminary set of 3D MC templates 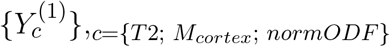 was generated by averaging of all registered volumes of T2w, cortex mask and normalized ODF channels (Fig. 2A).

#### 2.4.2 Generation of a age-specific multi-channel templates

At the second iteration (Fig. 2B), we used registration with T2w + cortex mask + normalized ODF channels (Sec. 2.3) to align all datasets to the multi-channel template (Sec. 2.4.1):

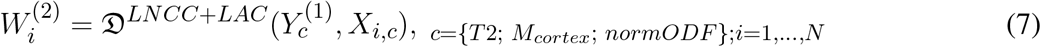

Next, the datasets were divided into 15 subsets according to PMA, to sample the range from 37 to 44 weeks PMA into 0.5 week time-windows. Each of the subsets *N^t^* contains 6-17 subjects depending on availability. The templates 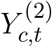 for each of the metrics (*c*) described in Section 2.2 were generated by robust weighted averaging the transformed metric maps *X_i,c_* in subsets *i ∈ N^t^*:

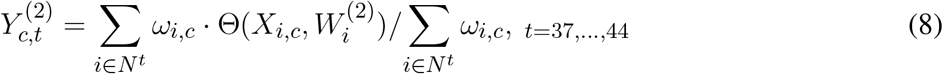

where Θ is the transformation operator, *c* is the list of all channels (See Fig. 2C). The voxel-wise weights *ω_i,c_* are binary maps with all values with > 1.5 standard deviations from the mean being excluded in order to minimise the impact of artifacts, cropped regions or misregistrations.

The templates 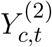 are biased towards 36 weeks reference space, we therefore calculate the transformations to remove this bias for each time-point. Because the registration is symmetric, we choose the inverse warps 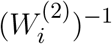 to create the transformation 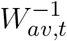 from the global reference space to the age-specific average space

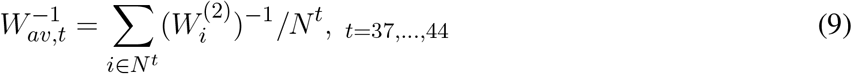

Similarly, we create average inverse affine transformation 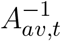 by selecting only the scaling and sheering components, followed by averaging and inverting.

#### 2.4.3 Parametrized 4D multi-channel atlas

In the final step we construct a continuous 4D spatio-temporal multi-channel model of the developing neonatal brain (Fig. 2C) by fitting the Gompertz growth curves to the time-dependent average metric maps and transformations. We propose the following form of the Gompertz function since it allows interpretation of both growth rate (*γ*) and peak in time (*τ*):

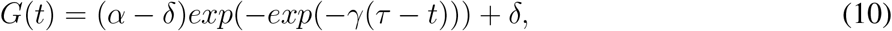

where *t* is the time point, *α* and *δ* control the upper and lower limits of *G*(*t*), *γ* represents the growth rate and *τ* is the center point corresponding to the growth peak. The model was fitted to the time-dependent average metric maps 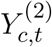 and transformations 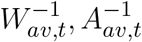 using least square minimisation to produce continuous spatio-temporal maps in reference space as well as average inverse transformations:

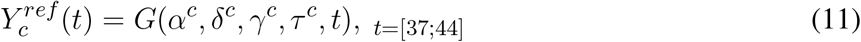

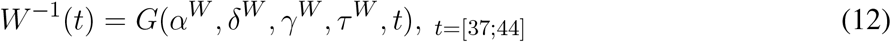

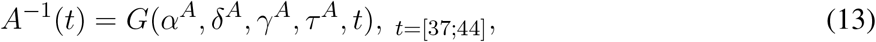

where *α^c^*, *δ^c^*, *γ^c^* and *τ^c^* are the interpretable Gompertz model parameters of metrics c = {T1w; T2w; T1w/T2w; mDWI; ODF: SH ODF, TDI; DTI: MD, RD, FA; DKI: MK; NODDI: ODI, FISO, ICVF; *μ*FA; Jacobian} and *t* is continuous over 37 to 44 weeks PMA range. Unbiased spatio-temporal maps *Y_c_*(*t*) are obtained by applying nonlinear transformation *W*^−1^(*t*) followed by affine transformation *A*^−1^(*t*) to the biased spatio-temporal maps 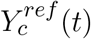.

### 2.5 Parcellation of WM regions

The dHCP structural atlas (Schuh et al., 2018) already provides parcellations of cortical and subcortical regions, we therefore specifically focus on WM tracts and transient regions. We first propagated the parcellation map of the major WM tract regions from M-CRIB-WM atlas (a single subject template at 41 weeks PMA, Alexander et al. (2020)) by registration to our T2w 41 week template 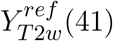.

Then we performed the MRTrix3 probabilistic tractography (Tournier et al., 2010) in *Y_ODF_* (41) channel for each of the 54 WM regions with propagated labels as seeds. This was followed by manual refinement of all labels in 3DSlicer (Fedorov et al., 2012) based on the thresholded TDI maps for individual tracts and the FA and T2 channels. The procedure was performed in three iterations. The output labels were stored in the atlas reference space resampled to 0.5 *mm* isotropic resolution to account for finer features.

The transient WM regions were localised as regions with high rates of signal changes during 37 to 44 weeks PMA. The parcellation was generated semi-automatically from the *γ^av^* map obtained by averaging growth rate *γ^c^* for T1w, T2w, RD and FISO channels. The *γ^av^* map was thresholded at experimentally selected value 0.25, followed by manual refinement. In addition, the parcellation map was masked with the dHCP atlas cortex mask in order to avoid inclusion of any regions affected by possible misregistrations in the cortex.

### 2.6 Atlas-based region-specific analysis

In order to assess the application of the proposed approach for the atlas-based region-specific analysis, we performed comparison of term and preterm cohorts. The analysis was based on both the WM and *γ^av^* parcellation maps. At first, all subject (selected 40 preterm and 140 term subjects scanned between 38 and 43 weeks PMA range) are registered to the PMA-matched atlas space (Sec. 2.3) with T2w, ODF, cortex and ventricle mask channels. It was identified experimentally, that adding the ventricle mask channel improves registration results for preterm subjects since preterm brains commonly have enlarged ventricles. Therefore, it was used for all subjects in the term-preterm comparison study. The average values for each of the structural and dMRI metrics were computed per each of the investigated regions using weighted averaging with only the values with the difference < 1.5 *st.dev* from the mean being included. Then, we assessed the association between the extracted metrics and the PMA at scan and the GA at birth using the ANOVA analysis. The output p-values were corrected for multiple comparisons.

### 2.7 Implementation details

Taking into account the differences in resolution between the input structural (0.5 *mm*) and diffusion datasets (1.5 *mm*), we chose 0.75 *mm* isotropic resolution as the optimal for the MC atlas. It also should be noted that unlike the previous dHCP atlas (Schuh et al., 2018), the proposed pipeline does not employ Laplacian sharpening of the atlas in order to avoid introducing artificial features into the interpretable maps.

The LAC metric for MC registration of ODF channels was implemented in MRtrix3 (Tournier et al., 2019). In addition, we implemented LNCC Demons metric (Avants et al., 2008) in MRtrix3 for registration of the structural channels. It was experimentally identified that multi-resolution 0.5; 0.75; 1.0 and SH order *l_max_* = {0; 2; 4} schemes and 3 voxel radius for the local neighbourhood for both structural and ODF channels produce high quality for deformable registration of the investigated datasets. We used the standard MRtrix3 regularisation of gradient update with 1 voxel standard deviation and displacement fields based on Gaussian smoothing with 0.75 voxel standard deviation. The proposed 4D GM fitting step (10) was implemented in MRtrix3. The ANOVA analysis for comparison between the term and preterm subjects was performed in RStudio (RStudio Team, 2020) using function `lm`.

## 3 RESULTS AND DISCUSSION

### 3.1 Multi-channel registration

We have previously demonstrated that the proposed MC registration improves overall alignment of cortical and WM regions when driven by both structural and ODF channels (Uus et al., 2020). Here we show that adding cortex mask *M_cortex_* channel further improves alignment of the cortex. We investigated three scenarios for the MC registration to the reference space (average 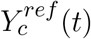 templates from the generated atlas): (I) *T*2*w* + *M_cortex_*, (II) *T*2*w* + *M_cortex_* + *FA*, (III) *T*2*w* + *M_cortex_* + *ODF* channels and (IV) *T*2*w* + *M_cortex_* + *ODFM_ventricles_* channels. We tested the performance on 12 term datasets from 42.00 to 42.57 weeks PMA range. The alingment was quantitatively assessed by NCC between transformed individual maps and templates *Y_c_*(*t*) of the corresponding PMA, using the TDI channel in the dilated WM region and T2w channel in the cortex and ventricles regions (highlighted in yellow in Fig.3A).

**Figure 3.**
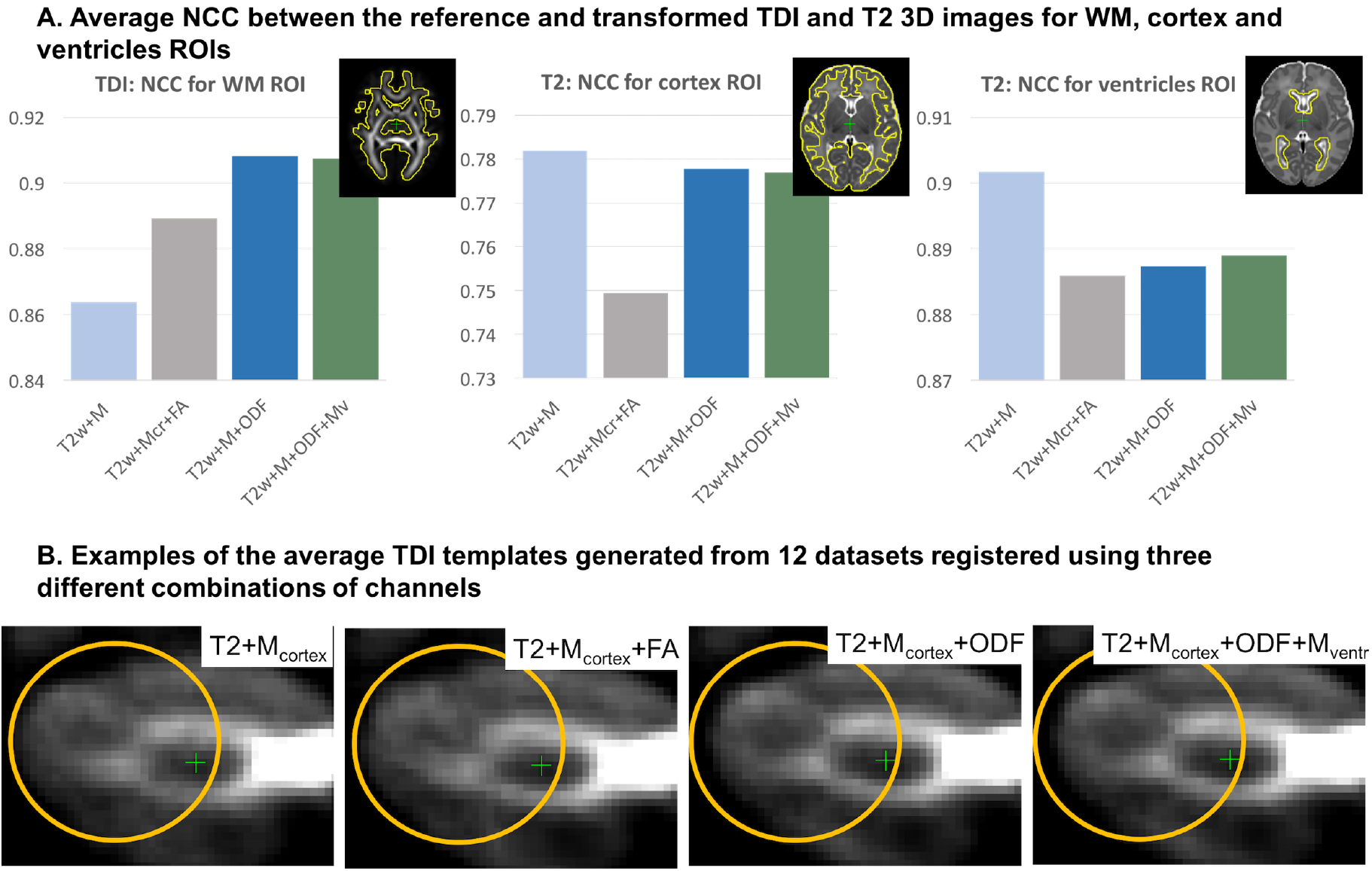
**A:** Comparison of MC registration results for three combinations of channels: *T*2*w* + *M_cortex_* (I,light blue), *T*2*w* + *M_cortex_* + *FA* (II,gray), *T*2 + *M_cortex_* + *ODF* (III,dark blue) and *T*2 + *M_cortex_* + *ODF* + *M_ventricles_* (IV,green). The performance was measured by NCC for the WM, cortex and ventricle regions between *Y_c_*(*t*) and transform TDI map and T2w image. The results are statistically significant with p < 0.001 for all cases apart from the combinations I and III for the cortex, II and III for ventricles and III and IV for all regions, which produced similar results. **B:** Examples of the average TDI templates in the cerebellum region generated from the datasets registered using different combinations of channels.

The *T*2*w* + *M_cortex_* + *ODF* MC registration (dark blue) produced the highest NCC within the WM for the aligned TDI maps. It also showed similar results to the structural only registration *T*2*w*+*M_cortex_* (light blue) within cortex region for the aligned T2 images. This was achieved by the additional cortex mask channel thus resolving the limitation reported in our previous work (Uus et al., 2020). The *T*2*w* + *M_cortex_* + *FA* MC registration showed significantly lower NCC potentially due impact of not well defined cortical features in FA map. Both *T*2*w* + *M_cortex_* + *ODF* and *T*2*w* + *M_cortex_* + *FA* (gray) combinations showed lower performance in the ventricle region. Addition of the ventricles mask only slightly improved the results in several cases. This was primarily due to the fact that the ventricles are quite small and the quality of the segmentation is not sufficient. At the same time, since the preterm subjects commonly have enlarged ventricles, we used it to improve the results for the preterm cohort study in the atlas-based analysis (Sec.2.6,Sec.3.4). The advantage of the *T*2*w* + *M_cortex_* + *ODF* MC registration is demonstrated visually by better defined average TDI map of the cerebellum shown in Fig.3B.

### 3.2 4D multi-channel atlas of normative neonatal brain development

The resulting multi-channel 4D atlas 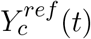 in the reference space (36 weeks PMA dHCP atlas (Schuh et al., 2018)) is given in Fig. 4. Unbiased atlas *Y_c_*(*t*) obtained after application of average inverse warps for 38, 41 and 44 weeks PMA time points is presented in Fig. 5. We can observe nonlinear changes due to cortical folding in the T2w templates. Volumetric expansion/contraction due to growth the is visible in the Jacobian maps.

**Figure 4.**
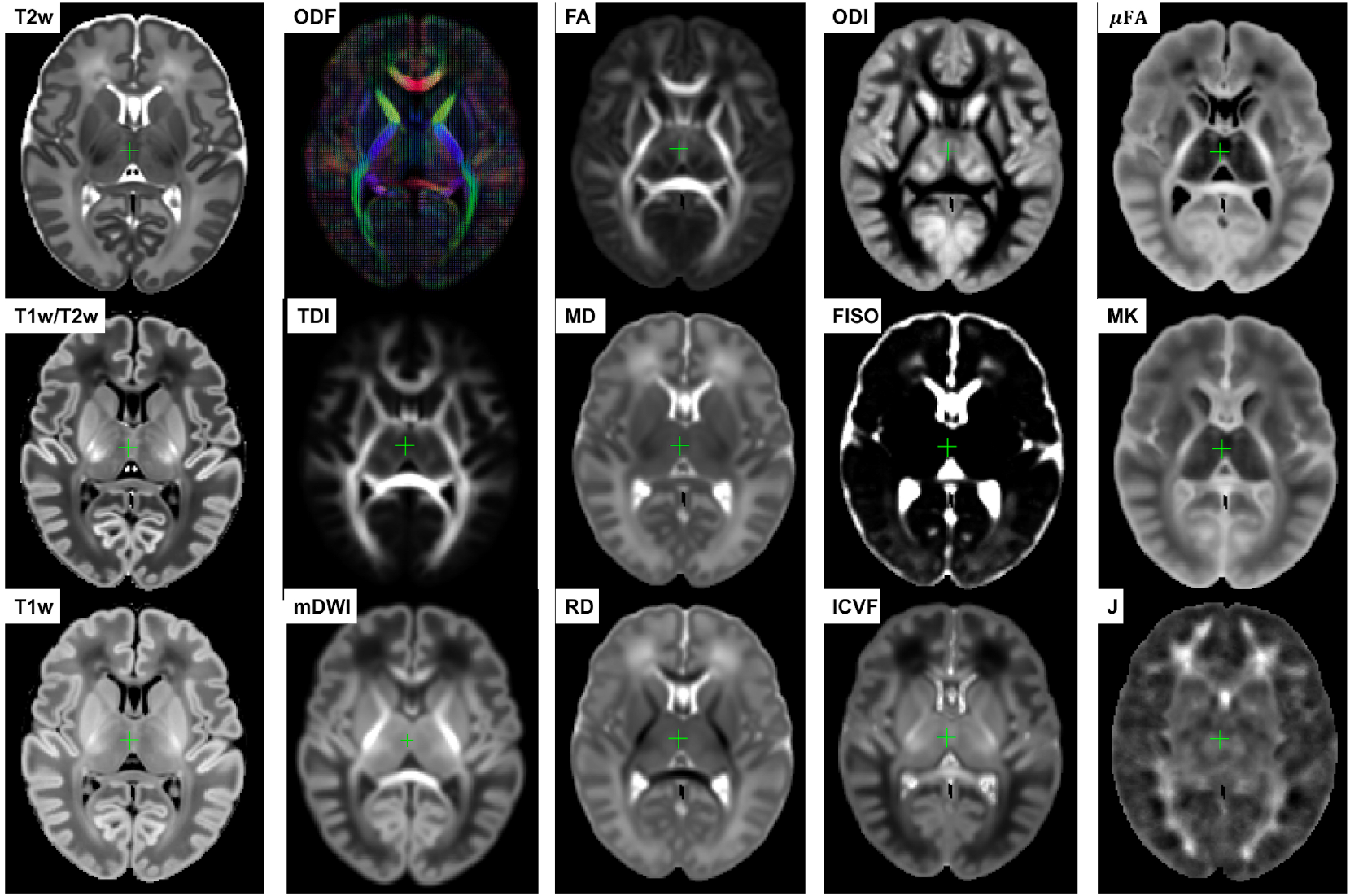
Multi-channel 4D atlas in the reference space (corresponding to 36 weeks PMA). Structural channels: T1, T2, T1/T2 and Jacobian; ODF channels: SH ODF, mDWI, TDI; DTI channels: MD, RD, FA; DKI channel: MK; NODDI channels: ODI, FISO, ICVF; *μ*FA.

**Figure 5.**
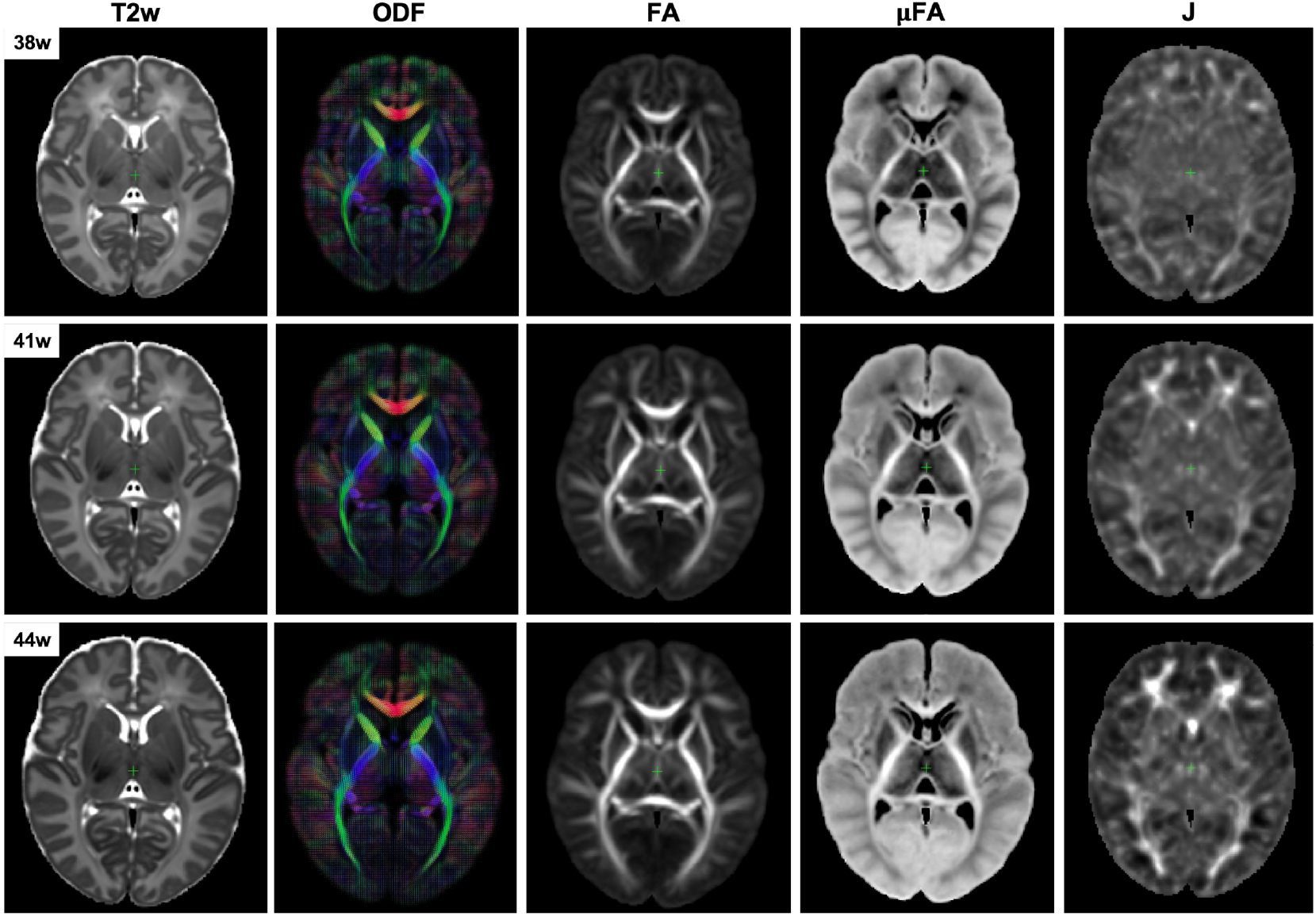
Example unbiased 4D atlas channels at 38, 41 and 44 weeks PMA. The corresponding Jacobian maps (J) are shown in the reference space.

The WM parcellations map with 54 ROIs created in the atlas reference space (Sec. 2.5) for the region-specific analysis of the metric values is shown in Fig. 6. The label annotation information follows the original annotations in Alexander et al. (2020).

**Figure 6.**
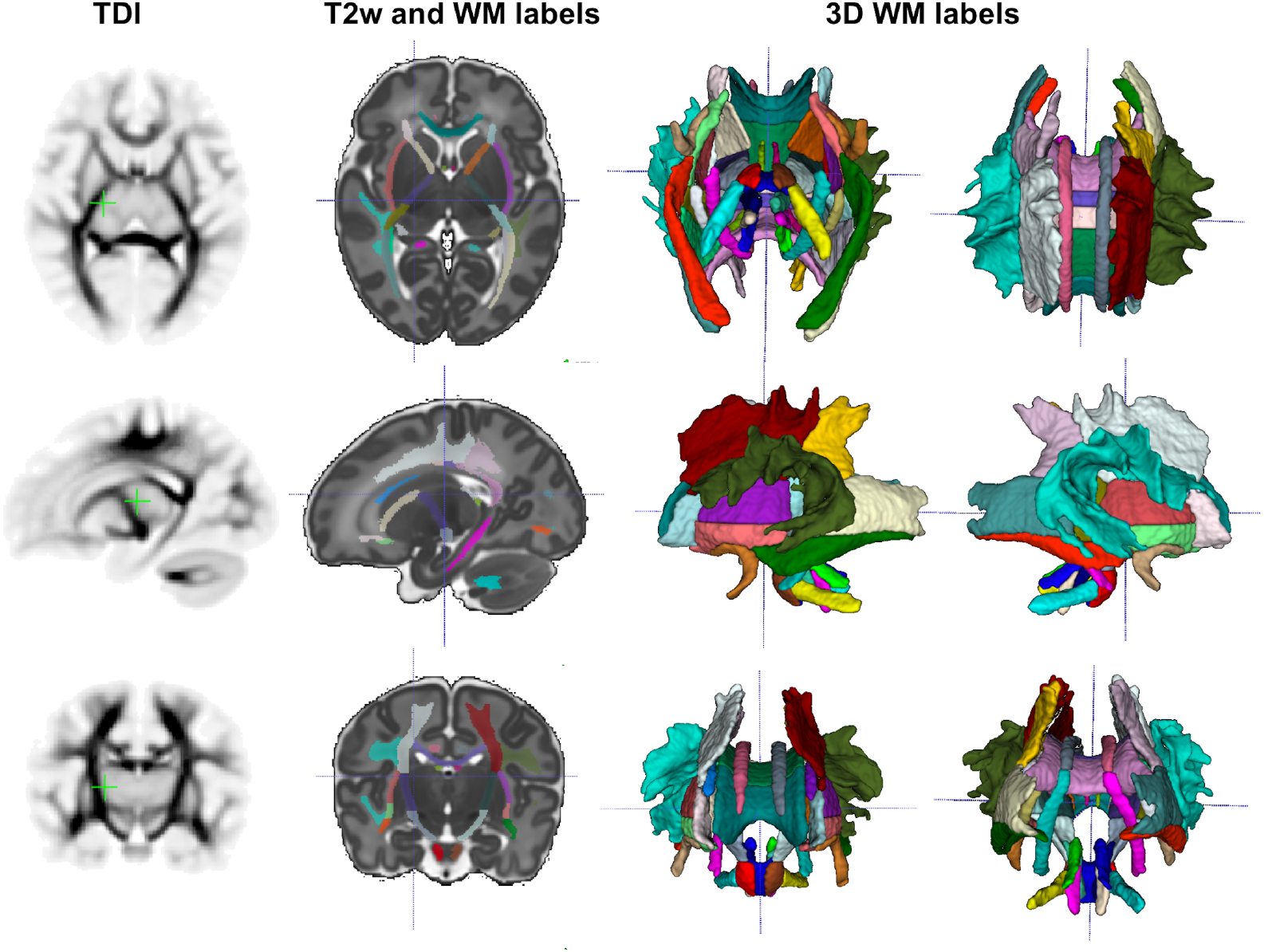
The WM parcellation map in the atlas reference space. The 54 ROIs are based on the structures defined in the M-CRIB-WM atlas (Alexander et al., 2020). The corresponding TDI map highlights the WM pathway regions.

Fig. 7.A presents the parcellation map of the transient regions identified by high rates of signal changes during 37 to 44 week PMA segmented from the average *γ^av^* map (Fig. 7.B). The parcellation map has 24 regions with the majority being consistent with the transient fetal compartment regions described in the recently introduced extended MRI scoring systems of cerebral maturation (Pittet et al., 2019) including periventricular crossroads, Von Monakow WM segments and subplate. We also identified fast developing regions within the cerebellum and subcortical grey matter.

**Figure 7.**
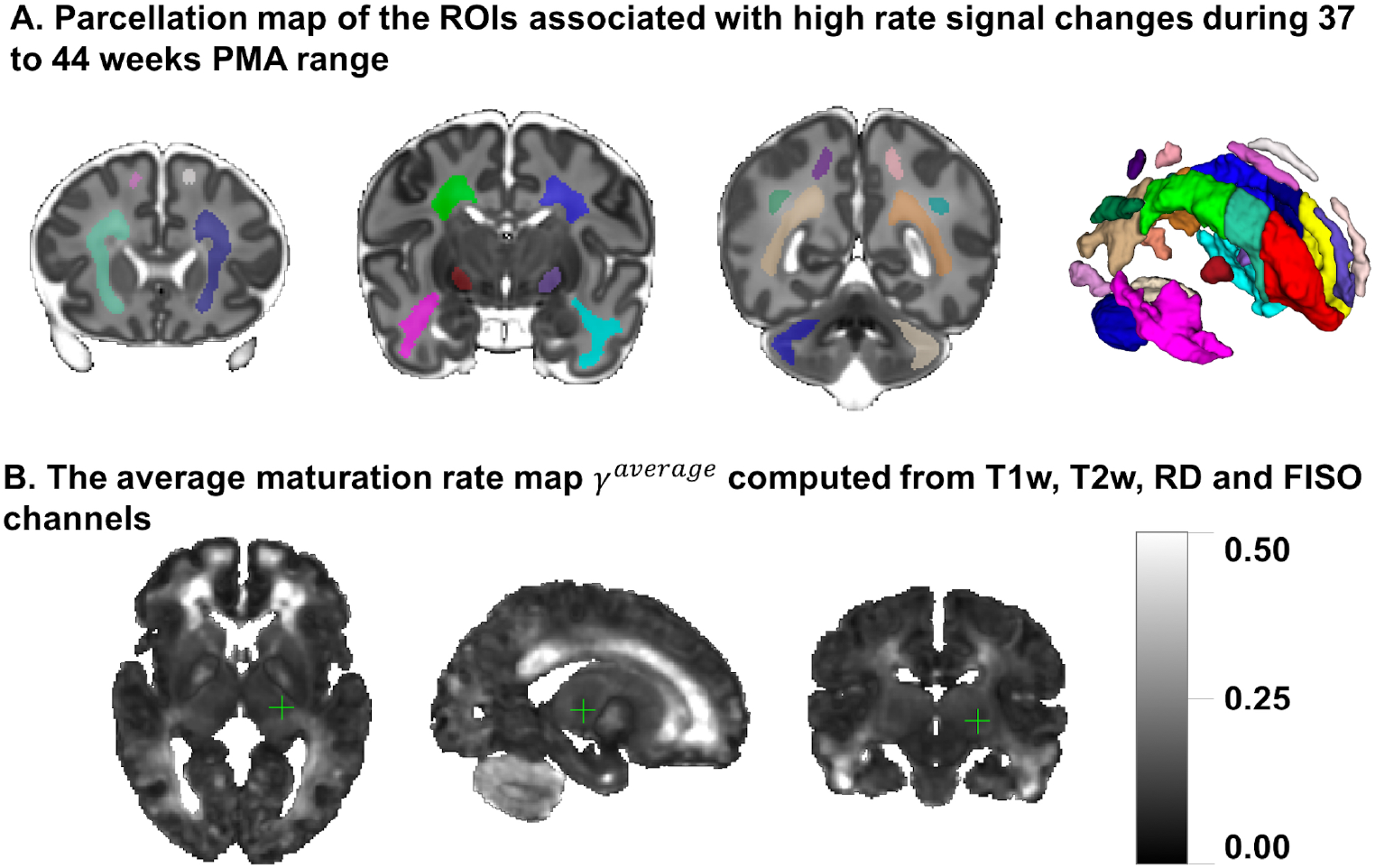
**A**: The parcellation map of 24 paired regions identified by high change rates during 37 to 44 week PMA. **B**: The average maturation rate map *γ^av^* computed from T1w, T2w, RD and FISO channels.

We also calculated *R*^2^ scores to evaluate the Gompertz model fit. Our results confirmed that GM offers higher *R*^2^ scores than linear regression with p<0.001 for the combined *γ* and WM parcellation map region. The primary regions where the GM fitting outperformed linear fitting corresponded to the *γ^av^* parcellation map (0.662∓0.189 vs. 0.652∓0.190 with p<0.001) with the high signal change rates. Examples of non-linear patterns are shown in Figures 9-11. Relatively small improvement in *R*^2^ however suggests, that linear fit offers reasonable approximation during this time-window.

### 3.3 Visual analysis of normal neonatal brain development

Fig. 8 shows probabilistic tractography generated from the ODF channel and the corresponding T1w channel (in the reference space) in the frontal WM region at 38, 41 and 44 weeks PMA time points. The increase in the T1w signal (known to be sensitive to proliferation of cells and myelin precursors and decreasing water content (Girard et al., 2012)) can be linked to the developing WM pathways seen in tractography (highlighted in red circle). The graphs show the corresponding increasing intensities in age-specific average templates 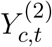 and fitted signal values 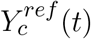 of the TDI and T1w channels computed in the small Von Monakow WM segment (Pittet et al., 2019) highlighted in yellow in the T1w channel.

**Figure 8.**
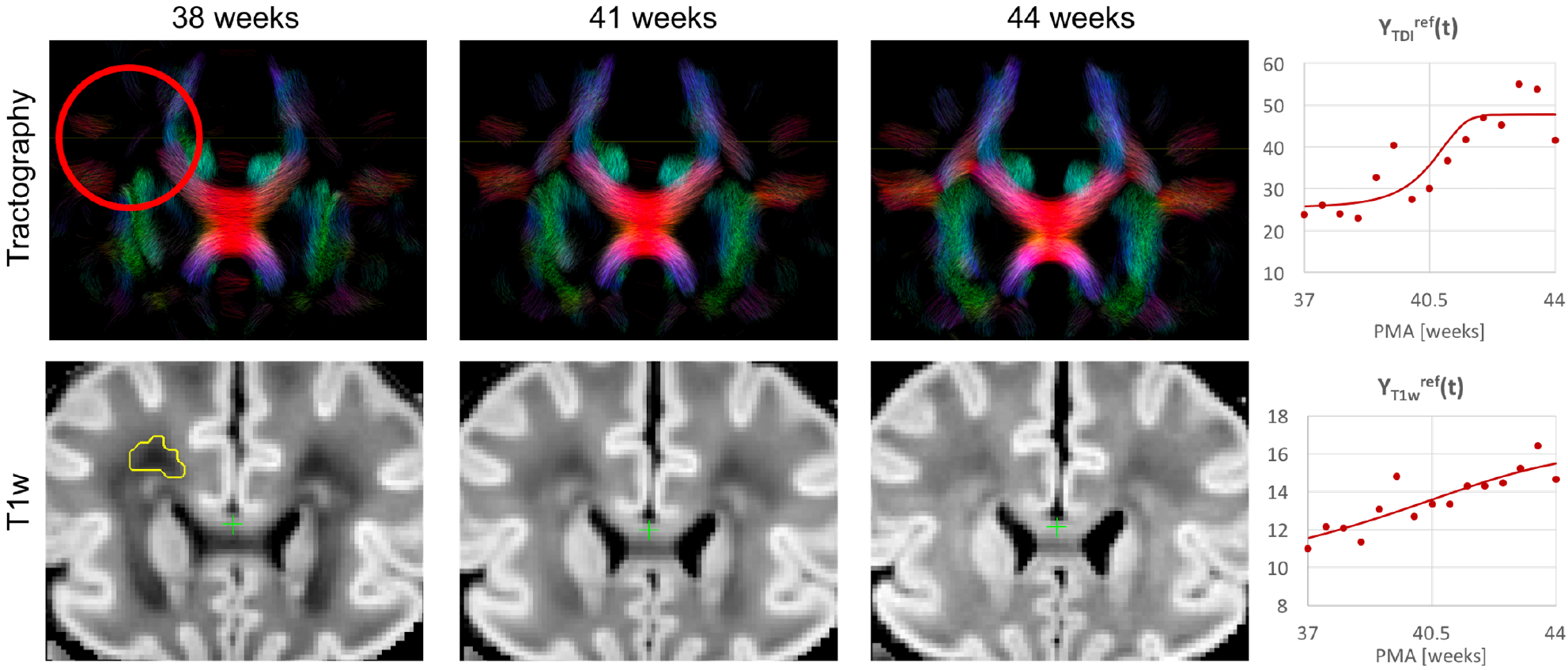
Whole brain probabilistic tractography generated from the ODF channel and the corresponding T1w channel (in the reference space) in the frontal WM region at 38, 41 and 44 PMA weeks time points. The developing WM pathway (red circle) can be linked to increasing T1w signal intensity (yellow region). The graphs show the signal in age-specific templates 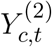 and fitted Gompertz model 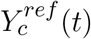 in the TDI and T1w channels averaged over the region highlighted in yellow.

Figures 9-11 show examples of the signal intensity changes in time and the corresponding growth rate maps *γ^c^*. The regions highlighted in yellow have significant growth peak offsets in time (≥0.2 weeks from the 40.5 weeks central time point) in *τ^c^*. The graphs show average signal values in 15 discrete age-specific templates 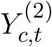 and the corresponding fitted signal 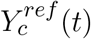 calculated within small 3×3×3 voxel regions at specific locations, including the right posterior limb of internal capsule (PLIC), superior corona radiata, periventricular crossroads, corpus callosum, Von Monakow WM segment and cerebellum.

**Figure 9.**
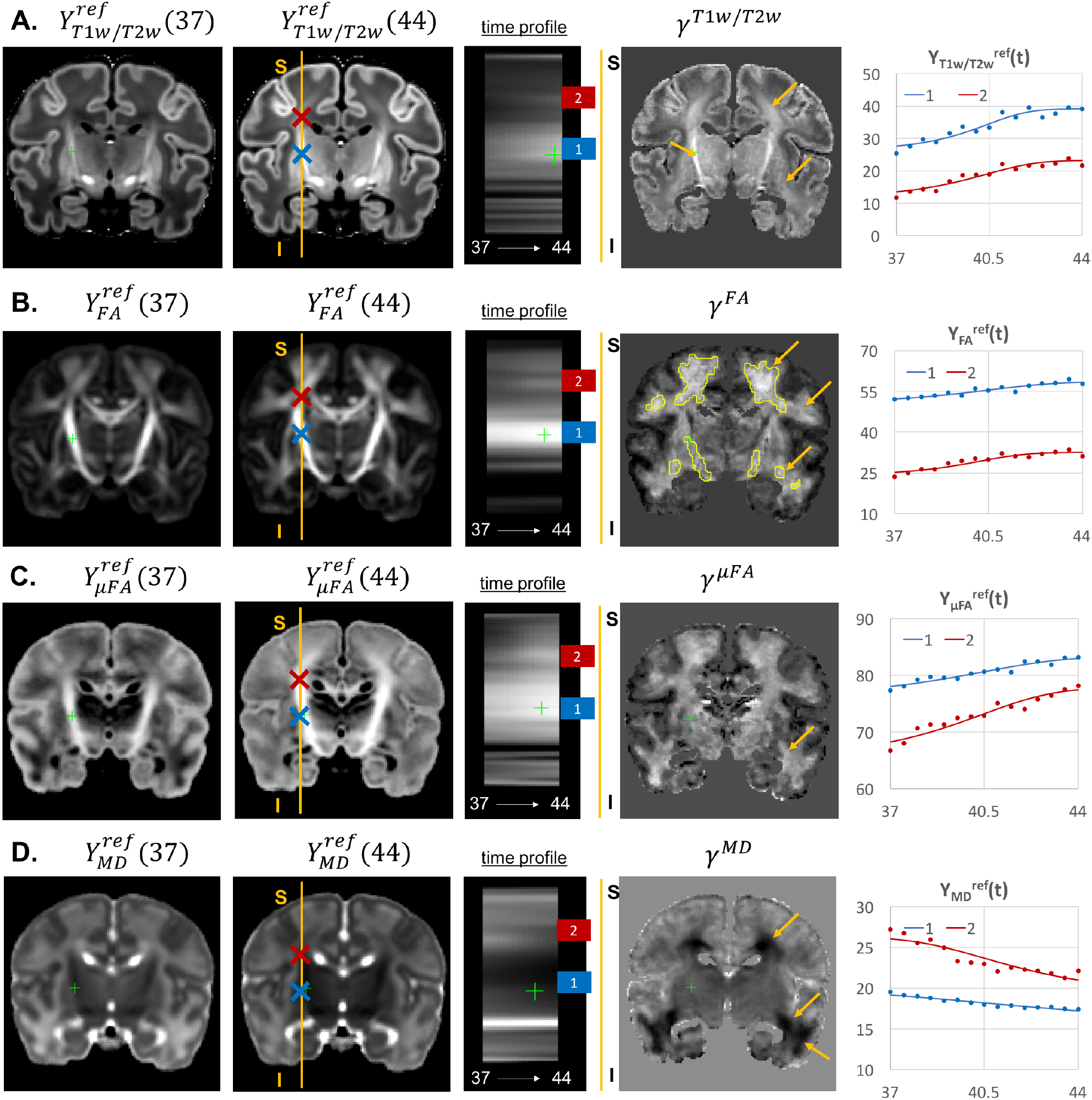
Examples of the signal changes in time (in the reference space) in T1w/T2w **(A)**, FA **(B)**, *μ*FA **(C)** and MD **(D)** channels. First column: 37 week template. Second column: 44 week template. Third column: signal change in time. Fourth column: *γ^c^* maps. Fifth column: Signal change in time in age-specific templates 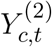 and fitted Gompertz model 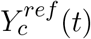 computed over 3×3×3 voxel regions in two locations: PLIC (blue) and superior corona radiata (red). The regions highlighted with yellow contours have significant growth peak offset in *τ^c^*.

The WM tracts are characterized by different maturation times and rates (Iida et al., 1995). The T1w/T2w contrast (linked to myelination by Glasser and Van Essen (2011)) shows signal increase from 37 to 44 weeks (Fig. 9.A). The *γ*^*T*1*w/T*2*w*^ map and the average signal graphs *Y*_*T*1*w/T*2*w*_(*t*) confirm that the rate of T1w/T2w signal increase is the highest in the PLIC region (blue) and the corona radiata (red). The *τ*^*T*1*w/T*2*w*^ parameter of the Gompertz model is 40.5 weeks in both regions which is agreement with the previously reported myelination milestones (Counsell et al., 2002; Wang et al., 2019). There is also a noticeable increase in the cortical T1w/T2w signal, also previously reported by Bozek et al. (2018), which may be due to myelination or the increased cell density (Girard et al., 2012). Both FA and *μ*FA signals (Fig.9.B-C) gradually increase in all WM regions in agreement with the trends reported in (Feng et al., 2019). The *μ*FA map shows generally higher degree of changes than FA, potentially due to the increasing crossing fiber effect, while in *γ^F A^*, the more prominent WM changes are observable primarily in the corona radiata, sagittal stratum and superior longitudinal fasciculus (highlighted with arrows). The *γ^MD^* map of the MD channel (Fig.9.D) shows high decrease in the superior corona radiata, sagittal stratum and the transient fetal compartments associated with WM maturation (Pittet et al., 2019) including the periventricular crossroads and subplate regions (highlighted with arrows). The MD signal is slowly decreasing the PLIC region as can be seen in the corresponding graph (blue). All of the presented *γ^c^* maps also show significant changes in the periventricular parietal crossroad regions (highlighted with arrows) with the significant decrease in MD and increasing in T1w/T2w.

Given the fixed number of streamlines used for probabilistic tractography, there is a notable redistribution of the TDI amplitude from the main to proximal WM tracts (Fig 10.A). The corresponding growth rate *γ^TDI^* map is positive in the frontal (anterior corona radiata) and thalamic radiation WM regions (highlighted with arrows) and negative in the internal capsule. The R-L time profile in the frontal region (Von Monakow WM segment, blue) shows the increased track density at 44 weeks. The average TDI signals 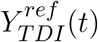 in this region (blue) and the corpus callosum (red) are also characterized by a significant degree of nonlinearity. NODDI FISO component (Fig. 10.B) shows a prominent reduction in the same frontal region which is in agreement with the expected decrease of water content and progressing maturation of WM pathways (Girard et al., 2012). Similarly to TDI, the average FISO signals 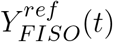 in the investigated WM ROIs have nonlinear shape with the steep decrease the occurring during the 39.5-43 weeks period. Similar decrease is observed in T2w signal (Fig. 10.C). The FISO channel in the sagittal view in Fig. 11.D also demonstrates similar patterns in the periventricular crossroads (red).

**Figure 10.**
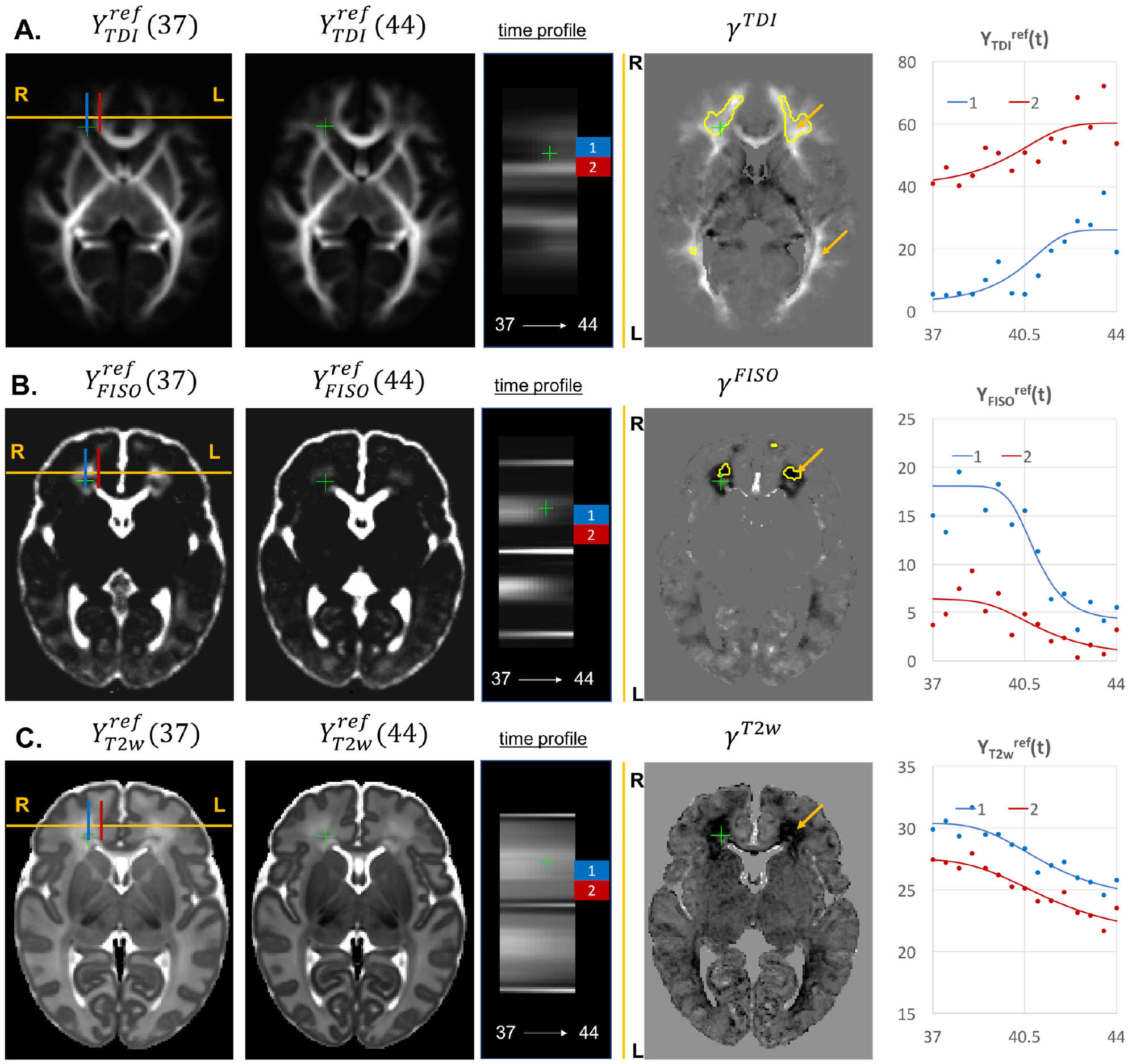
Examples of the signal changes in time (in the reference space) in TDI **(A)**, FISO **(B)** and T2w **(C)** channels. First column: 37 week template. Second column: 44 week template. Third column: signal change in time. Fourth column: *γ^c^* maps. Fifth column: Signal change in time in age-specific templates 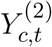 and fitted Gompertz model 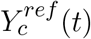 computed over 3×3×3 voxel regions in two locations: prefrontal corpus callosum (red) and Von Monakow WM segment (blue). The regions highlighted with yellow contours have significant growth peak offset in *τ^c^*.

**Figure 11.**
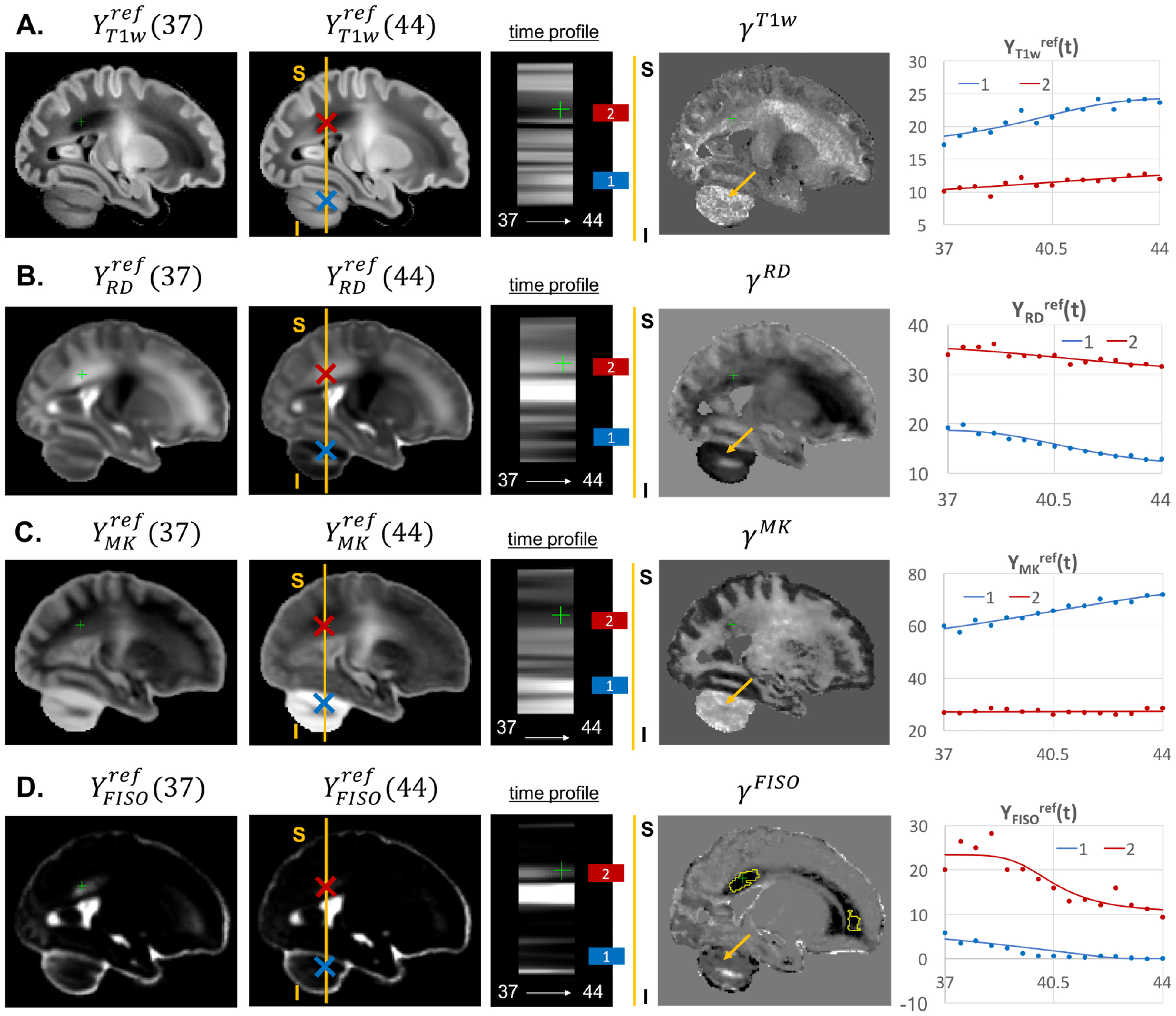
Examples of the signal changes in time (in the reference space) in T1w **(A)**, RD **(B)**, MK **(C)** and FISO **(D)** channels. First column: 37 week template. Second column: 44 week template. Third column: signal change in time. Fourth column: *γ^c^* maps. Fifth column: Signal change in time in age-specific templates 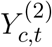 and fitted Gompertz model 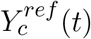 computed over 3×3×3 voxel regions in two locations:cerebellum (blue) and periventricular crossroads (red). The regions highlighted with yellow contours have significant growth peak offset in *τ^c^*.

Most of the channels also show prominent changes in the cerebellum associated with normal maturation (Fig.11, blue). The T1w signal intensity 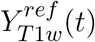 is gradually increasing due to WM maturation along with the increasing microstructural complexity reflected in the MK channel with the high *γ^MK^* map values and the expected decreasing trends of the RD 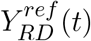 and FISO *Y_FISO_*(*t*) signals (potentially due to the decreasing amount of free water (Girard et al., 2012)).

### 3.4 Atlas-based region-specific analysis

We performed ANOVA analysis to assess influence of GA at birth on microstructure of WM regions delineated in our new atlas, with PMA as a cofounding variable. In order to assess the feasibility of using the ANOVA analysis for the investigated datasets, we performed linear fitting for each of the channels. The *γ^c^* values showed high correlation with the linear slope maps with the average NCC for all channels in the whole brain ROI 0.903∓0.085 (without CSF). This confirms that during 37 to 44 weeks PMA range linear approximation is acceptable.

Fig.12 visualises WM and transient regions in selected channels where average signal value was significantly associated with GA at birth. We found significant association of multiple indices with GA in the corona radiata, superior longitudinal fasciculus, corpus callosum and thalamic radiation. The T1w/T2w contrast was also significantly associated with GA in the internal and external capsules (Fig.12.A). We also found significant association of GA with *γ^av^* parcellation regions (Fig.12.B), which is in agreement with the prolonged existence of transient compartments in preterm subjects (Kostović and Judaš, 2006).

**Figure 12.**
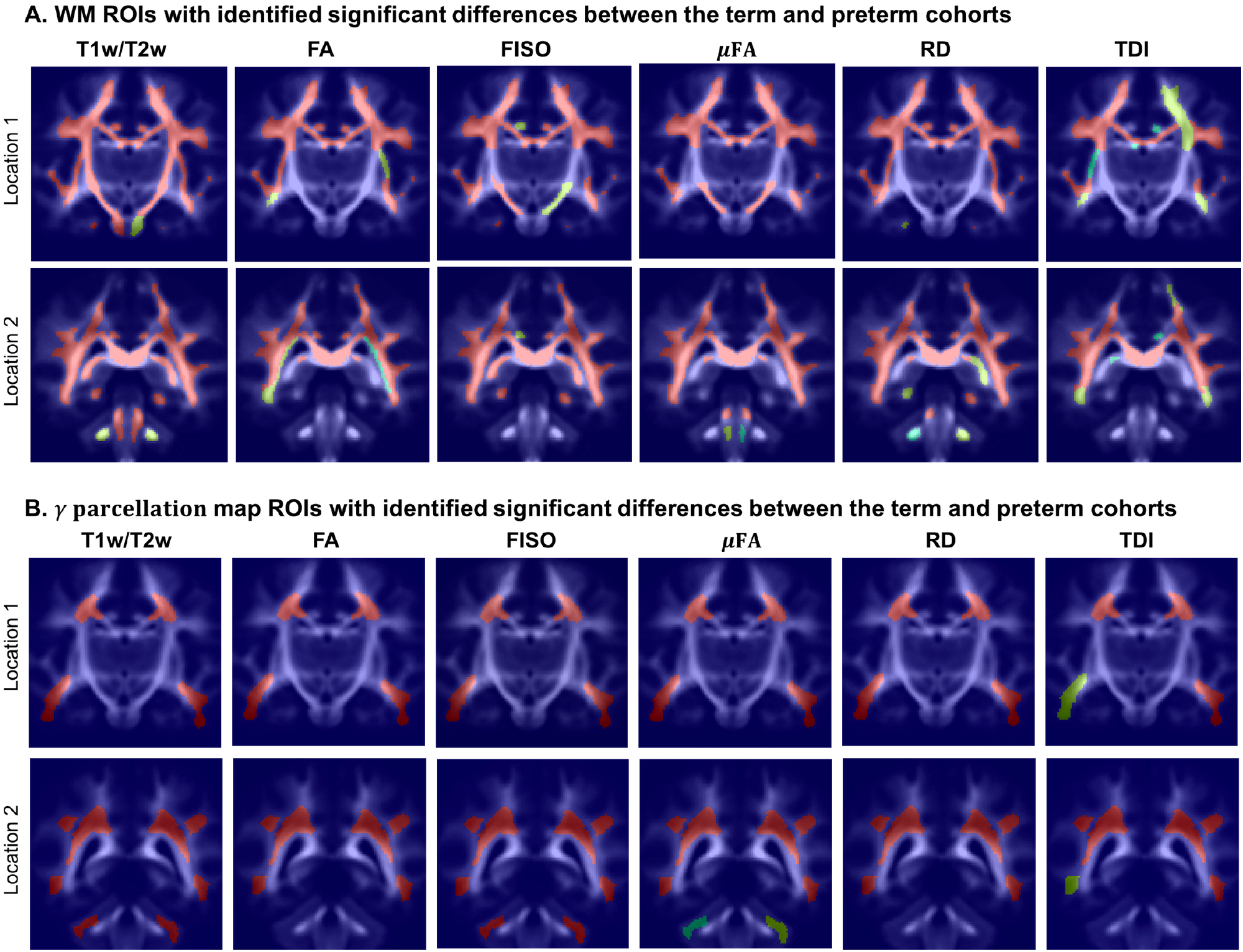
Atlas-based region-specific analysis. The regions significantly associated with GA at birth are highlighted with red (p < 0.001), yellow (p < 0.01) and green (p < 0.05) and overlaid over the averaged TDI map in two coronal view locations. **A**: WM parcellation regions. **B**: *γ^av^* parcellation regions.

Fig.13.A highlights differences in rate of maturation in *γ^c^* maps between term and preterm cohorts. The graphs in Fig.13.B show the average signal values in the frontal right Von Monakow WM segment (highlighted in yellow in the *γ^c^* maps). There is a clear increasing trend in T1w/T2w, FA and TDI for the term cohort along with the decreasing FISO and RD. However, the slopes for the preterm cohort are close to zero with high variance in the signal values. Furthermore, in this region, the preterm subjects are characterized by the significantly higher FISO and RD values and lower T1w/T2w, TDI and FA than the term cohort at the 42-43 week PMA period. This is consistent with the commonly reported lower FA and higher diffusivity values in preterm groups (Hermoye et al., 2006; Knight et al., 2018; Dimitrova et al., 2020), again suggesting delayed maturation of transient compartments in premature babies (Kostović and Judaš, 2006).

**Figure 13.**
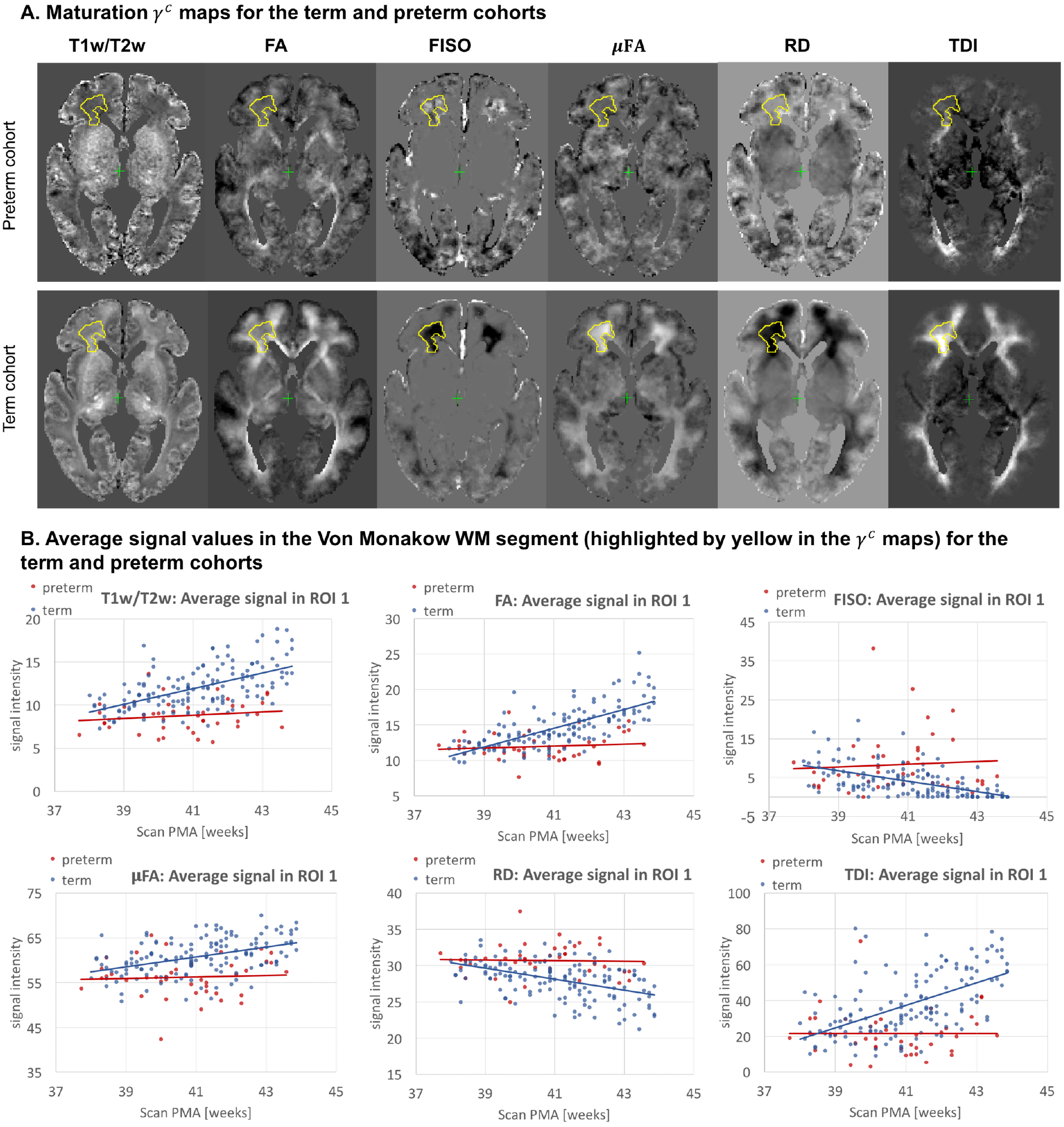
Atlas-based analysis: comparison of the term (140) and preterm cohorts (40) for 38 to 43 weeks scan PMA range for a subset of channels c= {T1w/T2w; TDI; RD; FA; FISO; *μ*FA}. **A**: The *γ^c^* maps of GM fitting for the term and preterm cohorts for 38 to 43 weeks PMA range. **B**: The mean signal values in the frontal WM ROI from the *γ^average^* parcellation map (highlighted in yellow in the gamma maps) for the term (blue) and preterm (red) cohorts for 38 to 43 weeks PMA range.

## 4 LIMITATIONS AND FUTURE WORK

The generated atlas is specific to dHCP acquisition protocols, which might limit application in terms of comparison with the datasets from other studies. However, the proposed tools can be easily applied to generate study- and acquisition-specific 4D MC atlases. We investigated relatively narrow neonatal period, and extension wider age range would improve reliability of Gompertz model fit and bring more insights into early brain development.

The study comparing term and preterm brain development included only 40 preterm subjects and they were not grouped with respect to specific types of anomalies, which can be addressed in future as more datasets become available. Including additional cortical and sub-cortical regions could also enrich the insights into normal and preterm microstructural brain development.

## 5 CONCLUSIONS

In this work, we proposed and implemented a novel pipeline for generation of continuous 4D multi-channel atlases. It is based on MC ODF+T2w guided registration and the Gompertz model fitting of both signal intensities and spatial transformations. The MC registration pipeline implemented in MRtrix3 employs the novel LAC similarity metric for ODF channels, LNCC metric for structural T2w and weighted fusion of the updates to the displacement fields. It also includes the cortex mask channel for better alignment of the cortex regions.

Based on the proposed methods, we generated a first continuous multi-channel atlas of the normal term neonatal brain development during 37 to 44 weeks PMA range generated from 170 subjects from dHCP project. The atlas contains 15 channels including structural (T1w, T2w and T1w/T2w contrast) and DWI-derived metrics based on ODF, DTI, DKI, *μ*FA and NODDI models. The GM fitting of the signal intensity and spatial transformation components in 4D allowed parametrization of the atlas. The output *γ* maps representing the rate of change can be used for interpretation of how maturation processes are manifested in different structural and diffusion MRI-derived metrics.

We also found that the fetal transient compartments (Pittet et al., 2019) have high contrast in the T2w, T1w, FISO, MD, RD and TDI *γ^c^* maps.

The atlas also includes detailed WM parcellation maps: (i) the map with the major WM tract ROIs based on the definitions from the recently introduced M-CRIB-WM neonatal atlas (Alexander et al., 2020) and (ii) the map of the regions associated with the high *γ* signal change rates during the normal WM maturation process. We tested the applicability of these parcellation maps for region-specific atlas-based studies on comparison between the term and preterm cohorts. We found significant effects linked to prematurity in the multiple WM regions including the transient fetal compartments. The atlas and the software tools are publicly available to support future studies of early brain development.

## AUTHOR CONTRIBUTIONS

A.U. prepared the manuscript, implemented the code for the extended MC registration, fitting and analysis, generated the 4D MC atlas and conducted the experiments. I.G. participated in implementation of the preprocessing and analysis code, the design of the study and interpretation of the results. M.P. developed the original code for SSD MC ODF registration in MRtrix3. B.D. performed preprocessing of the dHCP datasets. M.P., D.C. and J.D.T developed the tools for preprocessing and analysis of HARDI dHCP datasets. J.H., E.H., J.D.T, L.C-G. and A.P. developed MRI acquisition protocols for the neonatal dHCP datasets. L.C-G. developed the tools for preprocessing of structural dHCP datasets. J.V.H., A.D.E., S.C. and M.R are coordinators of the dHCP project. MD conceptualised the study and the methods, obtained the funding and supervised all stages of the research and preparation of the manuscript. All authors gave final approval for publication and agree to be held accountable for the work performed therein.

## ACKNOWLEDGMENTS

We thank everyone who was involved in acquisition and analysis of the datasets. We thank all participants and their families. The views expressed are those of the authors and not necessarily those of the NHS, the NIHR or the Department of Health.

## SUPPLEMENTAL DATA

## DATA AVAILABILITY STATEMENT

The generated MC atlas including all 4D fitted signal map channels, fitted 4D transformations, output GM fitting maturation map and both WM and *γ* parcellation maps together with the software tools used to generate the atlas will be available online after publication of the article.

The preprocessed datasets analyzed in this study will become available after the public release of the dHCP data.

## Footnotes

1 dHCP project: http://www.developingconnectome.org

2 dHCP weekly neonatal brain atlas: https://gin.g-node.org/BioMedIA/dhcp-volumetric-atlas-groupwise

